# Neuronal firing and waveform alterations through ictal recruitment in humans

**DOI:** 10.1101/2020.01.11.902817

**Authors:** Edward M. Merricks, Elliot H. Smith, Ronald G. Emerson, Lisa M. Bateman, Guy M. McKhann, Robert R. Goodman, Sameer A. Sheth, Bradley Greger, Paul A. House, Andrew J. Trevelyan, Catherine A. Schevon

**Affiliations:** Department of Neurology, Columbia University Medical Center, New York NY; Department of Neurosurgery, University of Utah, Salt Lake City UT; Department of Neurology, Weill Cornell Medical Center, New York, NY; Department of Neurosurgery, Columbia University Medical Center, New York NY; Department of Neurosurgery, Lenox Hill Hospital, New York NY; Department of Neurosurgery, Baylor College of Medicine, Houston TX; School of Biology and Health Systems Engineering, Arizona State University, Tempe AZ; Intermountain Healthcare, Murray UT; Institute of Neuroscience, Newcastle University, Newcastle upon Tyne, UK

## Abstract

Clinical analyses of neuronal activity during seizures, invariably using extracellular recordings, is greatly hindered by various phenomena that are well established in animal studies: changes in local ionic concentration, changes in ionic conductance, and intense, hypersynchronous firing. The first two alter the action potential waveform, whereas the third increases the “noise”; all three factors confound attempts to detect and classify single neurons (units). To address these analytical difficulties, we developed a novel template-matching based spike sorting method, which enabled identification of 1,239 single units in 27 patients with intractable focal epilepsy, that were tracked throughout multiple seizures. These new analyses showed continued neuronal firing through the ictal transition, which was defined as a transient period of intense tonic firing consistent with previous descriptions of the ictal wavefront. After the ictal transition, neurons displayed increased spike duration (*p* < 0.001) and reduced spike amplitude (*p* < 0.001), in keeping with prior animal studies; units in non-recruited territories, by contrast, showed more stable waveforms. All units returned to their pre-ictal waveforms after seizure termination. Waveshape changes were stereotyped across seizures within patients. Our analyses of single neuron firing patterns, at the ictal wavefront, showed widespread intense activation, and commonly involving marked waveshape alteration. We conclude that the distinction between tissue that has been recruited to the seizure versus non-recruited territories is evident at the level of single neurons, and that increased waveform duration and decreased waveform amplitude are hallmarks of seizure invasion that could be used as defining characteristics of local recruitment.

**Significance Statement:** Animal studies consistently show marked changes in action potential waveform during epileptic discharges, but acquiring similar evidence in humans has proved difficult. Assessing neuronal involvement in ictal events is pivotal to understanding seizure dynamics and in defining clinical localization of epileptic pathology. Using a novel method to track neuronal firing, we analyzed microelectrode array recordings of spontaneously occurring human seizures, and here report two dichotomous activity patterns. In cortex that is recruited to the seizure, neuronal firing rates increase and waveforms become longer in duration and shorter in amplitude, while penumbral tissue shows stable action potentials, in keeping with the “dual territory” model of seizure dynamics.

## Introduction

A complete understanding of the mechanisms underlying seizure pathology and dynamics depends on knowledge of the local neuronal activity, and what is driving that activity. Comparative animal models have long been used to gain insights into the underlying neuronal activity during seizures (Purpura *et al*., 1972; Fariello *et al*., 1976; Grone & Baraban, 2015), with the paroxysmal depolarizing shift (PDS) being regarded as the intracellular correlate of ictal discharges in animal models for more than half a century (Kandel & Spencer, 1961a, 1961b; Matsumoto & Marsan, 1964; Traub & Wong, 1982).

More recently, early PDSs have been shown to evolve into seizures *in vivo* (Steriade & Amzica, 1999), and PDSs have been recorded in resected human cortical tissue (Marcuccilli *et al*., 2010; Eissa *et al*., 2016). The PDS causes a decrease in action potential amplitude and an increase in half width – features that should impede standard spike sorting methods – and yet this phenomenon has not been reported in several studies of single unit activity during spontaneous human seizures (Wyler *et al*., 1982; Babb *et al*., 1987; Stead *et al*., 2010; Truccolo *et al*., 2011, 2014; Bower *et al*., 2012). In fact, beyond the PDS, altered action potential waveforms could be expected following recruitment of a recording site to a seizure due to alterations to Na^+^ and K^+^ concentrations in the intracellular and extracellular space or the effects of burst firing (Harris *et al*., 2000).

We have shown preliminary evidence of such potential alterations (Merricks *et al*., 2015). In tissue recruited to the seizure, traditional spike sorting methods can fail to cluster single units in human ictal recordings from neocortical layers 4/5, where neuronal cell body density is particularly high (Keller *et al*., 2018), thereby hindering the ability to track evidence of wave shape alterations or neuronal firing patterns during and after the ictal wavefront (Merricks *et al*., 2015). However, whether this originated from alterations to neurons’ intrinsic wave shapes or simply from interference of action potentials from nearby, highly active cells has been unclear.

Here, we present analyses of neuronal activity in the human brain during focal seizures using novel template matching methods in order to characterize action potential waveform alterations and single unit firing patterns, as the ictal wavefront approaches, recruits, and passes the local tissue. We hypothesize that, similar to observations in animal models, human focal seizures consistently display alteration of intrinsic action potential shapes upon ictal recruitment. Specifically, we hypothesize that the distinction between recruited and penumbral tissue is maintained at the level of single neurons, with recruited cells displaying reduced spike amplitude, and increased duration, an effect that is absent in penumbral sites demonstrating increased firing rates, but lacking typical seizure hallmarks.

## Materials and Methods

### Human recordings

Adult patients undergoing surgical evaluation for pharmacoresistant focal epilepsy at Columbia University Irving Medical Center (CUIMC) and University of Utah were implanted with either a 96 channel, 4 × 4 mm “Utah”-style microelectrode array (UMA; Blackrock Microsystems, Salt Lake City, UT) or Behnke-Fried style microwires (BF array; Ad-tech Medical Equipment Corp, Oak Creek, WI) simultaneous to standard clinical electrocorticography (ECoG) or stereo-electroencephalography (sEEG) respectively. UMAs were implanted into neocortical gyri based on presurgical estimation of the ictogenic region, with electrode tips reaching layer 4/5 (1.0 mm electrode length; layer confirmed via histology in Schevon *et al*. (2012)), while BF arrays consisted of 8 microwires protruding ~4 mm from the tips of clinical depth electrodes.

Neural data were recorded at a sampling rate of 30 kHz on each microelectrode with a range of ± 8 mV at 16-bit precision, with a 0.3 Hz to 7.5 kHz bandpass filter. ECoG and sEEG data were collected with a sampling rate of either 500 Hz or 2 kHz, with 24-bit precision and a bandpass filter of 0.5 Hz to ¼ the sampling rate. In UMAs, the reference was either subdural or epidural, chosen based on recording quality. In BF arrays, the reference was the ninth microwire within the bundle.

All procedures were approved by the Institutional Review Boards of CUIMC and University of Utah, and all patients provided informed consent prior to surgery. Clinical determination of seizure onset zone (SOZ) and seizure spread was made by the treating physicians and confirmed prior to analysis by two board-certified neurologists (CAS & LMB). All analyses were performed offline using custom scripts and toolboxes written in MATLAB (MathWorks, Natick MA). Code is available at https://github.com/edmerix.

The timing of the passage of the ictal wavefront at individual electrodes was calculated based on the MUA firing rate. A Gaussian kernel of 500 ms duration was convolved with the timings of all detected spikes in the MUA, and a sustained, significant increase in the resultant instantaneous firing rate was determined as the moment of local recruitment to the seizure (Smith *et al*., 2016). A sustained (> 1 s), significant increase had to be present for classification as ictal recruitment in order to discount single discharges or herald spikes. Ictal recordings without this signature of tonic to clonic MUA firing were determined to be penumbral. Note that unlike the SOZ, ictal recruitment and the penumbra are spatiotemporally dynamic. As such, a single location, unless at the true origin of the ictal activity, may receive the synaptic input of the upstream ictal activity but remain penumbral due to feed-forward inhibition initially, prior to the transition to the ictal state which may occur at any point during the electroclinical seizure event.

### Peri-ictal single unit discrimination

Initial spike sorting was performed on the peri-ictal period as per Merricks *et al*. (2015). Briefly, neural signals were symmetrically bandpass filtered between 300 Hz and 5 kHz (1000^th^ order FIR1) to extract multi-unit activity (MUA), from which extracellular action potential spikes were detected using a voltage threshold of 4.5 σ, where 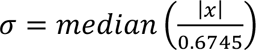, and *x* is the MUA from that channel. This method avoids the biasing effect of large spikes on channels with units with high firing rates (Quian Quiroga *et al*., 2004). Ictal periods were blanked so that spike sorting was only performed on stable spikes from the peri-ictal period.

Matrices of waveforms from each channel were created from 0.6 ms prior to 1 ms post each detection, and principal component based semi-automatic cluster cutting was performed using a modified version of the “UltraMegaSort2000” MATLAB toolbox (Hill *et al*., 2011). Artifactually large waveforms were removed by calculating the FFT on spikes up-sampled by a factor of 4, and removing those with power > 5 SD above the mean in frequencies above 2.5 kHz or below 500 Hz. Spikes removed in this manner were visually inspected to ensure correct classification as artefact. Clusters that satisfied the following criteria were accepted: (i) clean separation from all other clusters in the Fisher’s linear discriminant in principal component space; (ii) less than 1% contamination of the 2 ms absolute refractory period; (iii) no clear outliers based on the anticipated chi-squared distribution of Mahalanobis distances; and (iv) less than 1% of estimated false negatives as estimated by the amount of a Gaussian fit to the detected voltages fell below the threshold for detection, as described in Hill *et al*. (2011).

### Template matching through seizures

Ictal recruitment has been shown to impede standard spike sorting due to either interference of hypersynchronous activity, intrinsic waveform alterations, or both (Merricks *et al*., 2015; Fig. 1). We therefore developed novel methods in order to match waveforms from the ictal period, regardless of recruitment, to their putative neuronal source based on templates derived from the peri-ictal units. In contrast to standard spike sorting methods, these minimized false negatives at the expense of increasing false positives so as to avoid missing potential matches. Cluster boundaries were defined as the 3-dimensional convex hull surrounding the features in principal component space of the previously defined units from both the pre- and post-ictal period (Fig. 2). This method accounts for two situations: that neurons within recruited cortex maintain their wave shape but are obscured by interference from other nearby cells; or that there are occasional or consistent alterations to a neuron’s intrinsic waveform that are minor enough to be maintained within the convex hull of feature space. The convex hull allows for alterations to wave shape in any dimension (any direction away from the cluster’s centroid).

**Fig. 1.**
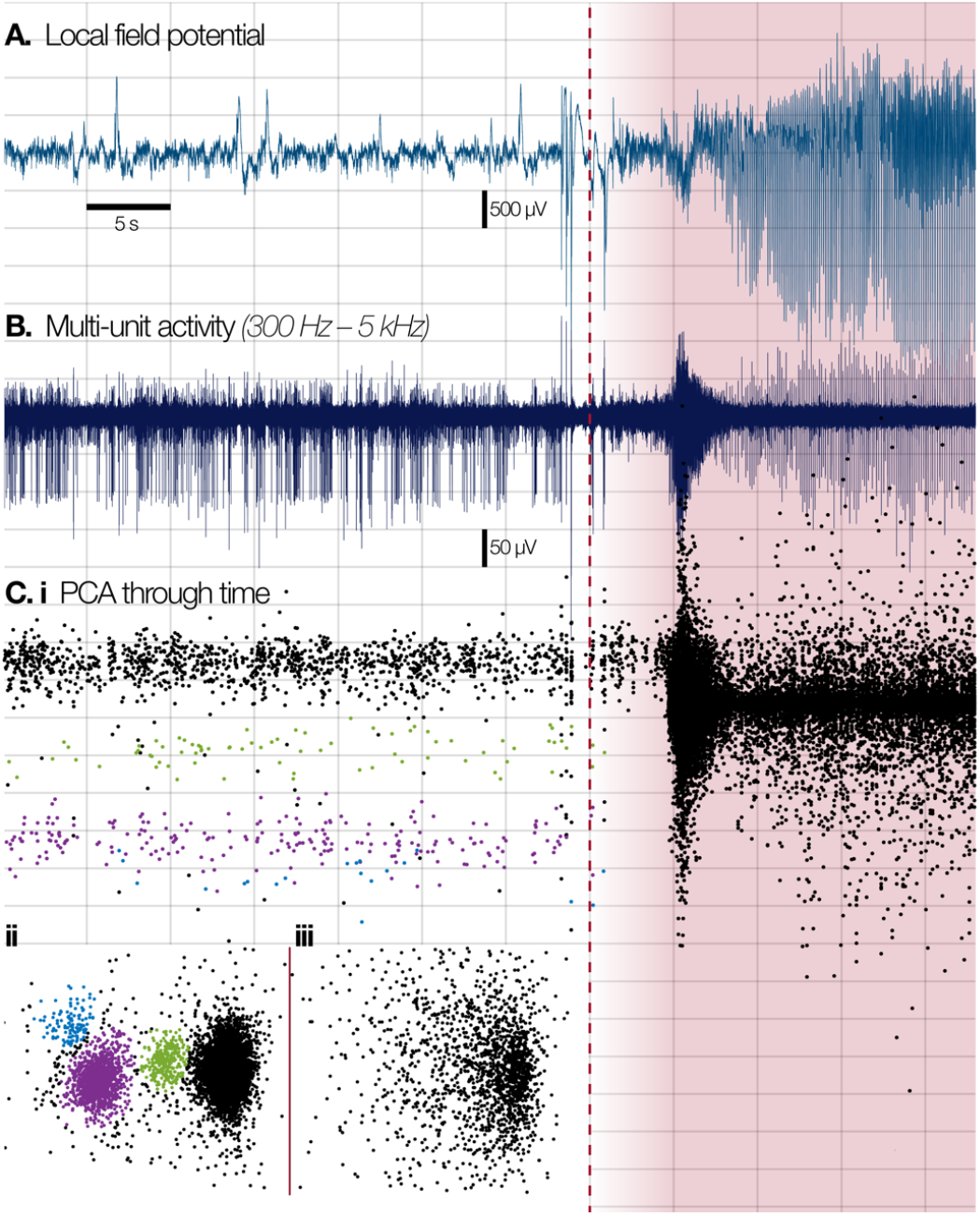
Effects of ictal recruitment on traditional spike sorting methods. Spike sorting relies on stable waveforms from nearby neurons, but ictal activity disrupts features used to cluster single units. **A.** Example broadband LFP from a single channel of a Utah array implanted in the posterior temporal lobe of a patient with pharmacoresistant epilepsy (Patient 4, seizure 1). Dashed red line denotes “global” seizure onset. **B.** Bandpass filtered signal between 300 Hz and 5 kHz of the same signal in A, showing stable single unit activity in the preictal period. **C.** First principal component score versus time (**i**) of all detected spikes in the multi-unit activity shown in B, with three clearly separable clusters highlighted, along with the multi-unit cluster from background distal cells (black). (**ii**) Equivalent first versus second principal component scores from the preictal period, and (**iii**), during the seizure. Note loss of well-defined clusters in principal component space.

**Fig. 2.**
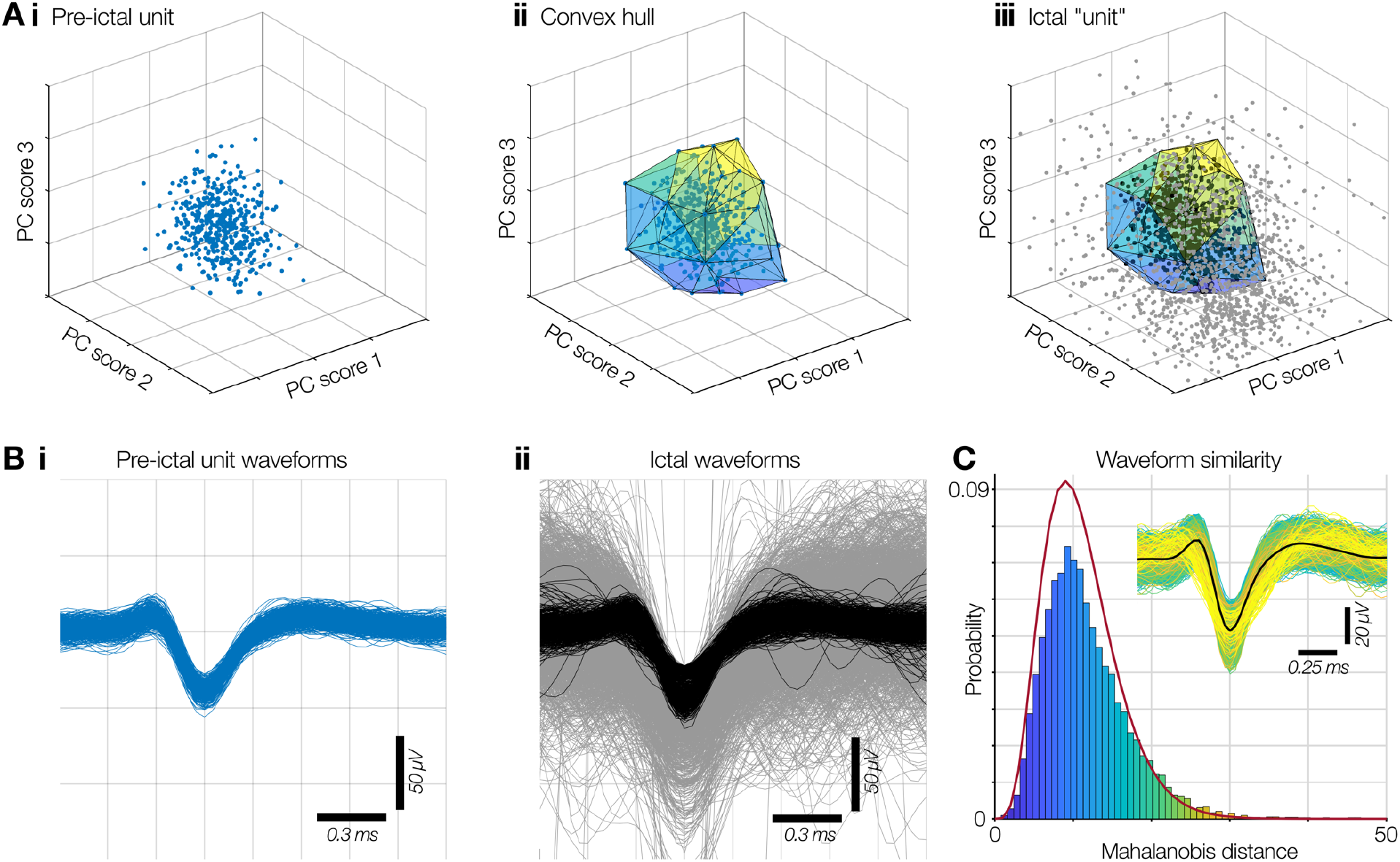
Template match spike sorting via convex hulls. **A.** Well-isolated neurons form distinct clusters in principal component space during interictal time points (**i**), around which a convex hull can be fitted to define boundaries in 3D space within which spikes that match that unit should exist (**ii**). Despite a lack of defined clusters during ictal activity in recruited cortex, this convex hull can be used to select waveforms that are likely to correspond to the preictal unit (**iii**; black), amongst distributed noise (grey). **B(i)** Waveforms from the preictal cluster shown in A(i). **(ii)** Waveforms matched using the convex hull shown in A(iii) (black) from the large distribution of spikes during a seizure (grey). **C.** The probability of each matched waveform originating from the same neuron as its preictal counterpart is calculated by calculating their Mahalanobis distance from the preictal cluster using the first *n* principal components that explain ≥ 95% of the variance. Outliers in the chi-squared distribution for *n* degrees of freedom denote likely incorrect matches.

To avoid ictal results being biased through differing methods, template matching was performed on spikes that were extracted from a period from 10 minutes prior to 10 minutes post seizures, including the ictal activity that had been blanked in the original peri-ictal spike sorting. Channels with unstable units during interictal periods were excluded. Units with no spikes in either the preictal or ictal time period after template matching and artefact removal were excluded from further analyses (*n* = 77).

Principal component scores were calculated on these spikes based on the previously defined principal components, and spikes that occurred within a peri-ictal unit’s convex hull were assigned to that cell. Mahalanobis distances were calculated for all matches, between their location in principal component space and all peri-ictal waveforms from that unit, on the first *n* principal components that explained > 95% of the variance in the data set (Fig. 2C). The expected distribution of Mahalanobis distances was calculated as the chi-squared probability distribution with *n* degrees of freedom. Spikes that had < 0.1% chance of occurring in the chi-squared distribution were excluded.

### Spike metrics

The full-width at half maximum (FWHM) was calculated by up-sampling each spike by a factor of 4, normalizing the spike voltages to between [-1, 1], and finding the difference between the zero-crossings either side of the spike’s trough. When calculating spike amplitude changes through the ictal transition, only units whose mean voltage at detection was at least 2.5 SD away from that channel’s threshold for detection were used, to minimize the floor effect from small units that dropped below threshold.

The probability that each spike arose from its assigned peri-ictal unit was calculated by fitting a separate Gaussian curve (with a maximum amplitude of 1) to the distribution of voltages at each data point in the original unit, and calculating the mean probability across all time samples. As such, a waveform passing through the most likely voltage at each separate time point for that unit would have a match probability of 1 (Fig. 3). Instantaneous firing rates were calculated by convolving the spike times with a Gaussian kernel (200 ms SD) with the amplitude scaled to that spike’s probability of matching the original unit, thereby creating probabilistic firing rates for each unit through time, by probability of when the spike occurred and probability that the spike originated from the putative neuronal source. As such, a waveform that had an average probability of 20% across all fitted Gaussians from each data point would contribute only 0.2 spikes s^−1^ at its most likely time point, while an exact match to the most likely voltage at each time point would contribute 1 spike s^−1^. Thresholds for significant increases and decreases in firing rate were calculated as 3 times the square root of the firing rate divided by the duration of the epoch, which approximates 3 SD for a Poisson distribution.

**Fig. 3.**
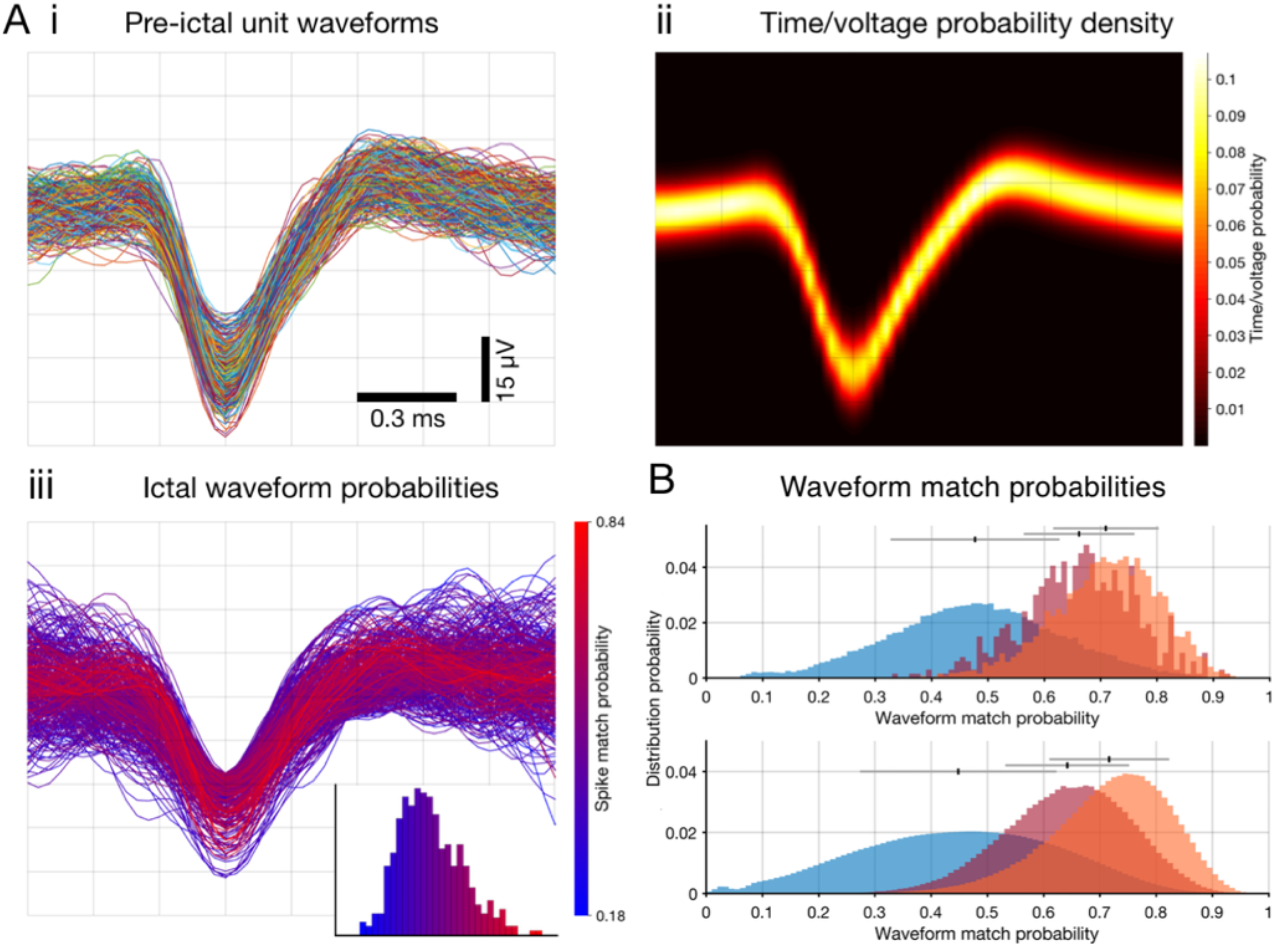
Waveform match probabilities. **A(i)**. All waveforms from a traditionally spike sorted unit in a 10 minute preictal epoch. The probability distribution of each voltage at each time point for these waveforms is shown in **(ii)**, and the spike match probabilities for ictal waveforms matching this unit are shown in **(iii)**, with the distribution of these probabilities inset. **B.** Upper panel shows the probability distribution of all matched waveforms (red), compared to the probability distribution for the original waveforms in the preictal time point (orange) for an example template-matched single unit. A bootstrap estimate of waveform matches expected by chance by comparing against waveforms from other electrodes is shown in blue. The lower panel shows the equivalent distributions across the full population. Mean ± SD is shown above the distributions.

Timing of FWHM alteration relative to the earliest ictal activity was determined by the earliest timepoint during the seizure that the mean FWHM remained above the preictal mean plus the preictal standard deviation for at least 1 second, calculated in a sliding window of 5 s duration with a time step of 50 ms, discarding windows with fewer than 5 spikes.

All statistical tests for significance were performed using the Mann-Whitney U test unless otherwise noted, due to the non-Gaussian distributions of data requiring non-parametric testing. For all tests, the level for statistical significance (*α*) was set to 0.05, and Holm-Bonferroni correction was applied in all instances of multiple tests.

## Results

We analysed ictal recordings from 27 patients undergoing invasive EEG monitoring as part of the presurgical evaluation for intractable focal epilepsy (Tables 1 & 2; age range = 19 to 55; 13 female, 14 male). Six patients were implanted with Utah microelectrode arrays (UMA; Blackrock Microsystems Inc, Salt Lake City, UT), and the remaining 21 patients were implanted with between 1 and 4 Behnke-Fried depth arrays with incorporated microwire bundles (BF arrays; Ad-tech Medical Equipment Corp, Oak Creek, WI). A total of 41 seizures were reviewed (10 UMA; 31 BF array), of which 27 demonstrated ictal recruitment through MUA firing rate calculation in UMAs, or subsequent waveform alterations in BF arrays (see Methods; UMA: 6 seizures from 3 patients; BF arrays: 21 seizures from 13 patients).

**Table 1.** Demographics and data for patients implanted with Utah arrays. Individual patient demographics for grid/Utah array implant cases, including the number of isolated single units found for each seizure in each patient. FIA: focal with impaired awareness; FTBTC: focal to bilateral tonic-clonic.

| Patient | Demographics | Implant location | Seizure type | Seizure onset | Seizures analyzed | # units |
| --- | --- | --- | --- | --- | --- | --- |
| 1 | 30 M | Frontal<br>(Left frontal convexity) | FIA | Left supplementary motor area | 1 | 79 |
| 2 | 39 M | Frontal<br>(Left dorsolateral frontal lobe) | FIA | Left frontal operculum | 2 | [61, 61] |
| 3 | 25 F | Temporal<br>(Left inferior temporal gyrus) | FIA | Left basal/<br>anterior temporal | 3 | [111, 101, 91] |
| 4 | 19 F | Temporal<br>(Right posterior temporal) | FTBTC | Right posterior lateral temporal | 1 | 110 |
| 5 | 24 M | Temporal<br>(Left inferior temporal gyrus) | FIA | Left mesial temporal with<br>spread to lateral temporal | 2 | [120, 133] |
| 6 | 30 M | Temporal<br>(Right mesial temporal gyrus) | FA | Right subtemporal | 1 | 240 |

### Dual activity types at seizure onset

To assess the presence of physiological spike shape changes across populations of neurons, template matching using convex hulls (Fig. 2, see Methods) was employed on each UMA-recorded seizure (Table 1; 1,107 single units in total; range of units per seizure: 61 – 240; mean ± SD maximum units per patient: 122 ± 63). The convex hull fits a 3-dimensional region around the peri-ictal unit’s cluster in principal component space, within which waveforms are assigned as a putative match to that neuron. This allows for assigning unit identities without the need for maintained cluster boundaries in the data, while accepting waveform alterations that alter the principal component scores. In total, 938 of the 1,107 units recorded on UMAs were successfully tracked through seizures using this method.

Individually, results from the template matching method in recordings from recruited tissue (see Methods; Patients 3, 4 & 5) showed decreases in spike amplitude and increases in spike full-width at half maximum (FWHM; Fig. 4, ictal waveforms in red), and were stereotyped across seizures within patient (Fig. 5). At the population level, units in recruited cortex displayed a significant global increase in FWHM (Fig. 6A; pre-ictal vs. ictal mean ± SD: 0.470 ± 0.137 ms vs. 0.611 ± 0.194 ms), with 457 (81.5%) of 561 single units showing a significant (*p* < 0.05) increase in FWHM during the seizure (Holm-Bonferroni corrected Mann-Whitney U test; range across seizures: 79% – 97%; see Table 3). Meanwhile, units in penumbral cortex showed only a minor increase in FWHM at the population level (Fig. 6B; pre-ictal vs. ictal mean ± SD: 0.414 ± 0.009 ms vs. 0.429 ± 0.099 ms) with only 9 (5.8%) of 156 single units showing a significant (*p* < 0.05) increase in FWHM during the seizure (Holm-Bonferroni corrected Mann-Whitney U test; range across seizures: 4% – 16%; see Table 3). In a single case (Patient 6), the UMA was at the edge of the clinically defined ictal spread, and in this patient 105 (47.5%) of 221 units showed a significant increase in FWHM (pre-ictal vs. ictal mean ± SD: 0.408 ± 0.111 ms vs. 0.421 ± 0.152 ms). These spike shape alterations co-existed with stable spike shapes elsewhere on the UMA at the same time (Fig. 7), and this patient was incorporated into the penumbral dataset for population representation in the figures (Fig. 6). FWHM increases in recruited tissue were significantly larger than those in penumbral/edge case recordings (*p* < 0.001; one-tailed Mann-Whitney U test).

**Fig. 4.**
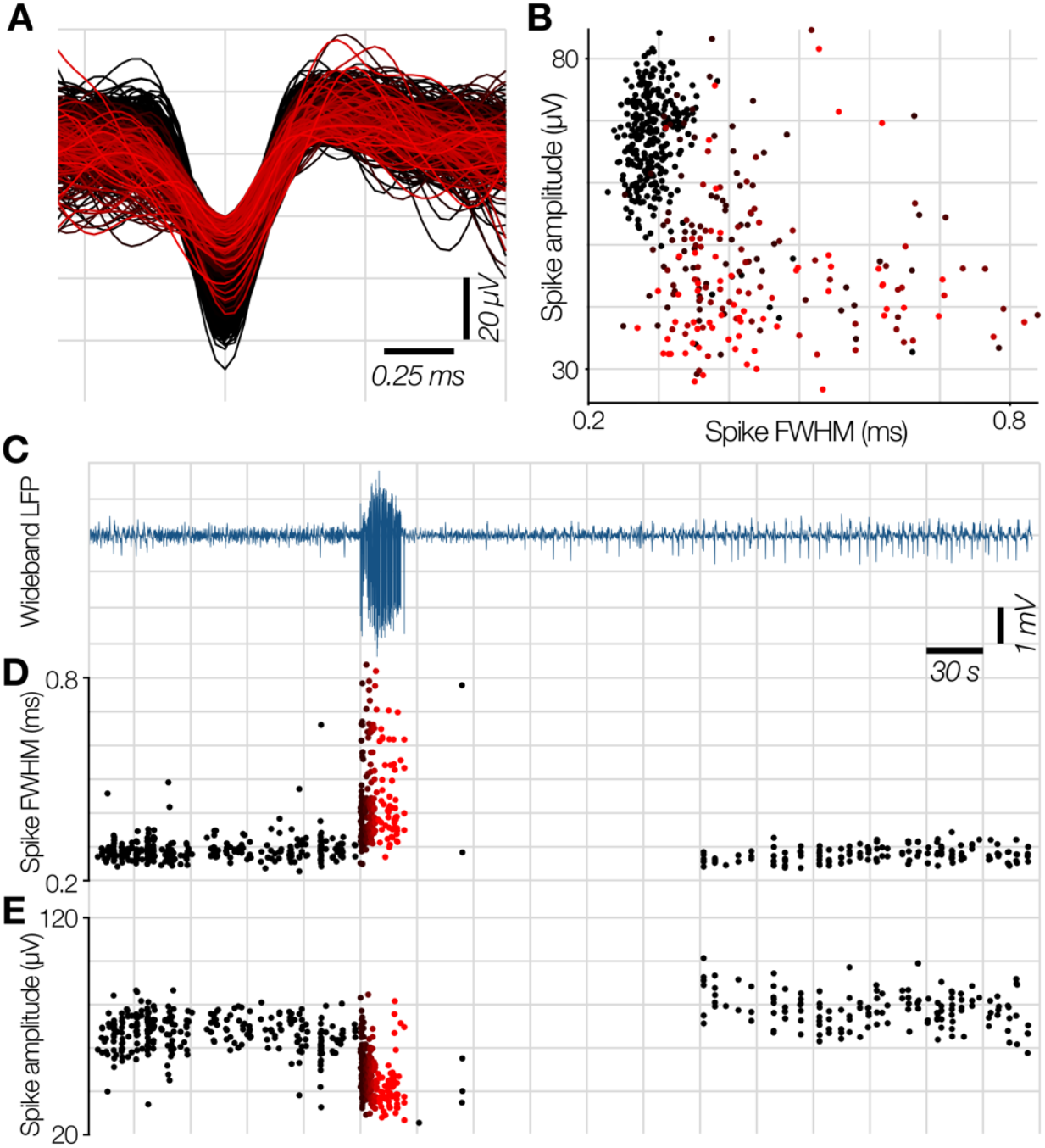
Example wave shape changes at ictal recruitment in template matched data. **A.** Waveforms from a convex hull-matched unit in Patient 5, seizure 2, showing reduction in amplitude and increase in FWHM during the seizure (red shaded waveforms, fading from black to red through the seizure; color maintained throughout figure). **B.** Spike amplitude versus FWHM in the unit in A, showing equivalent relationship to the high amplitude BF single unit in Fig. 8D. Wideband LFP from this channel through time (**C**), with time-locked FWHM (**D**) and amplitude (**E**) showing temporal relationship of spike shape changes through the seizure. Note the return to preictal values after seizure termination, and the lack of changes towards decreased FWHM or increased amplitude during the seizure despite the convex hull being equally permissive of alterations in any direction.

**Fig. 5.**
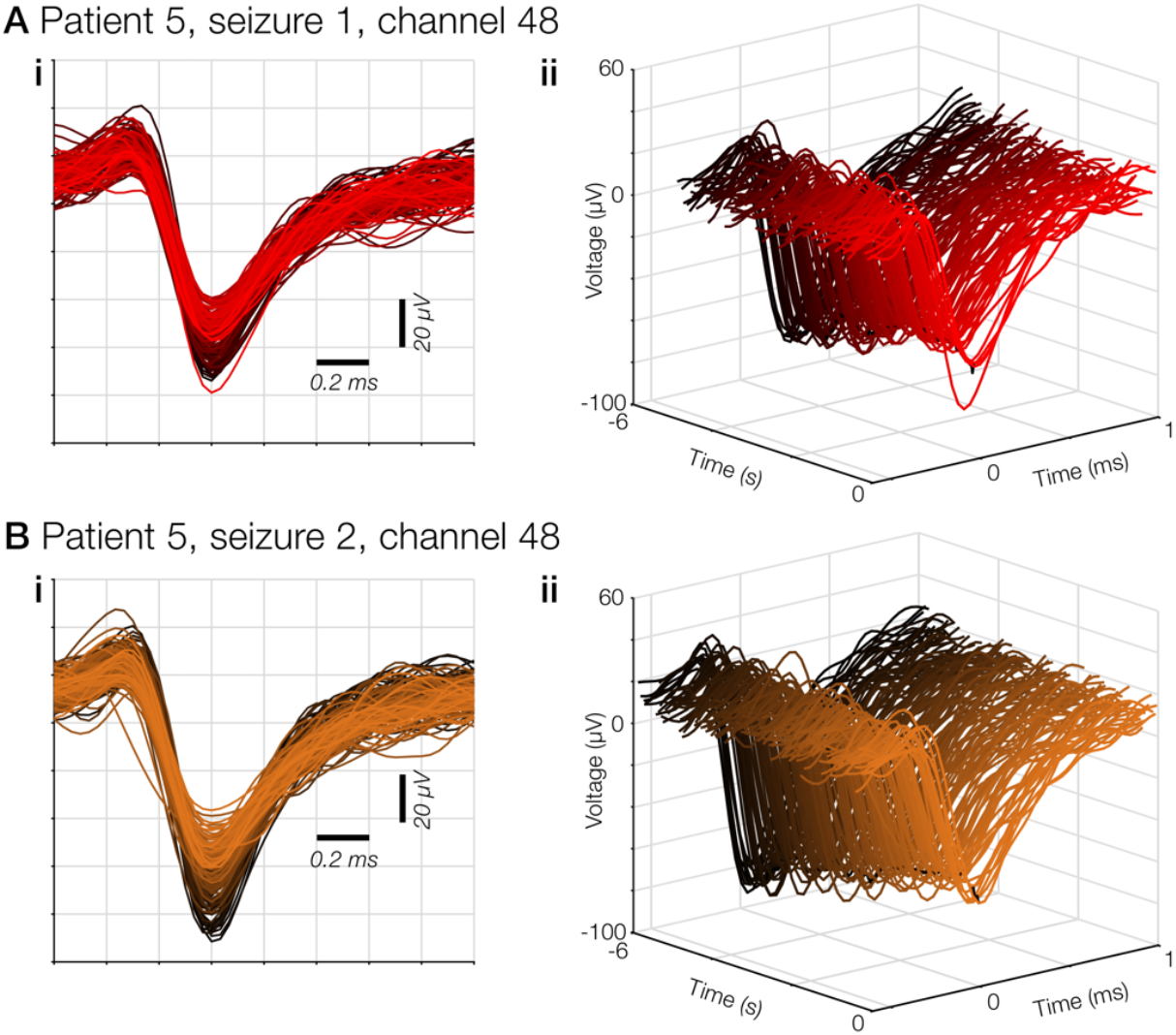
Stereotypy of waveform changes within neurons across seizures. Example waveforms from a single unit in patient 5 showing stereotypy of response of extracellularly recorded action potentials in the peri-recruitment period in 2 seizures separated by 22 hours (**A** and **B** respectively). All waveforms from the 6 seconds prior to ictal recruitment are shown overlaid in **(i)**, and plotted relative to time in **(ii)**, scales maintained throughout. Saturation of color fades from black at −6 seconds, to brightest at the moment of maximal firing rate of MUA at that electrode.

**Fig. 6.**
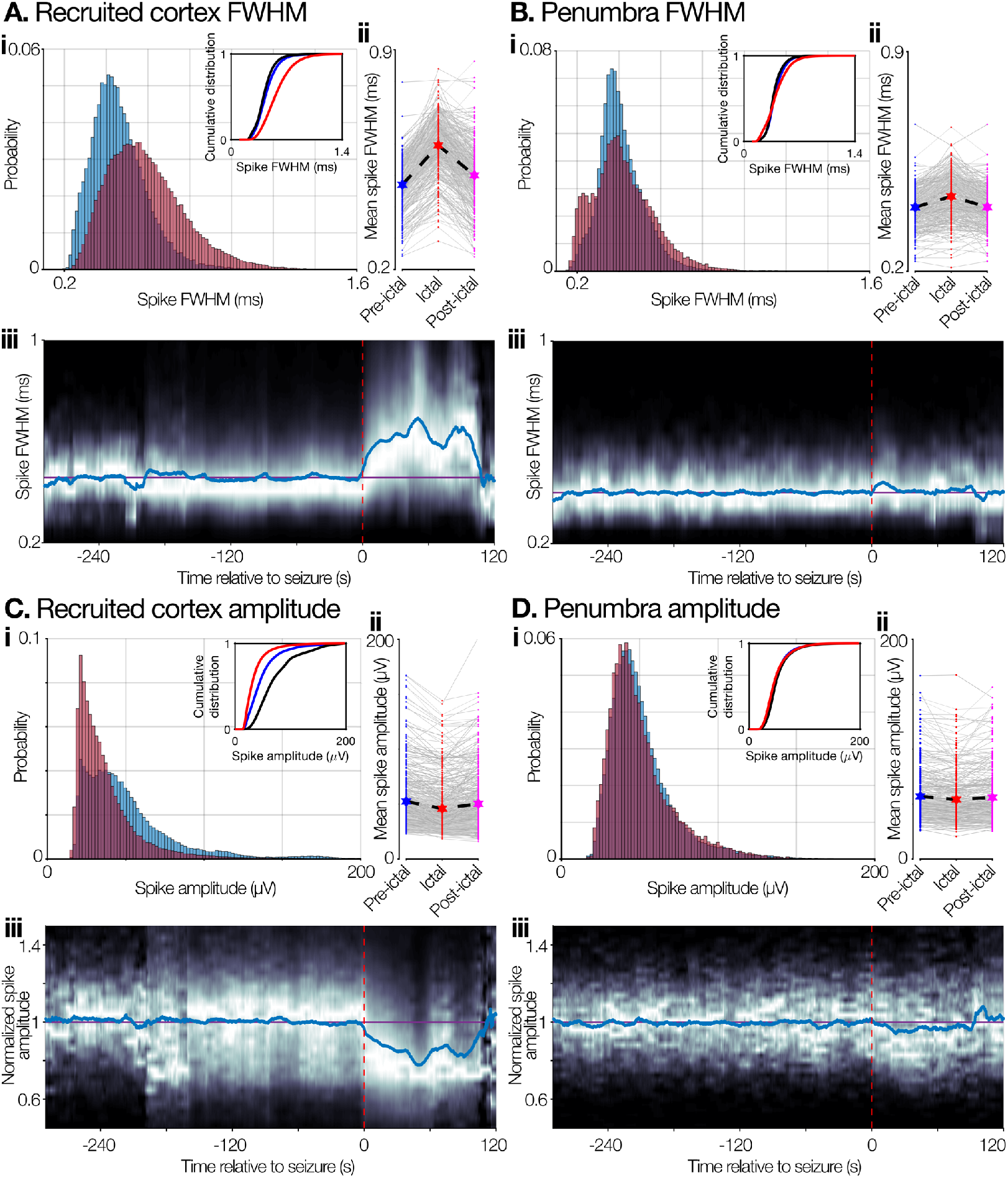
Population spike shape alterations in recruited cortex versus penumbral territories. **A & B (i)** Probability density plots of spike FWHM (full-width at half maximum) for every detected waveform in the preictal (blue) and ictal time periods for all seizures in recruited cortex (A; *n* = 625 units from 6 seizures in 3 patients) and penumbral tissue (B; *n* = 405 units from 4 seizures in 3 patients). Cumulative probability densities show same calculation on preictal, original data (black), **(ii)** Paired mean FWHM for each unit in the preictal (blue), ictal (red), and postictal (pink) epochs. Note return to preictal ranges after seizure termination. **(iii)** Spike FWHM of the population through time (10 second window, sliding every 100 ms). Brighter indicates more density, blue line shows the mean through time, purple line shows the mean value in the preictal period, red dashed line denotes “global” seizure onset. **C & D.** Same format as A & B, showing spike amplitude in place of FWHM, for recruited cortex and penumbra respectively.

**Fig. 7.**
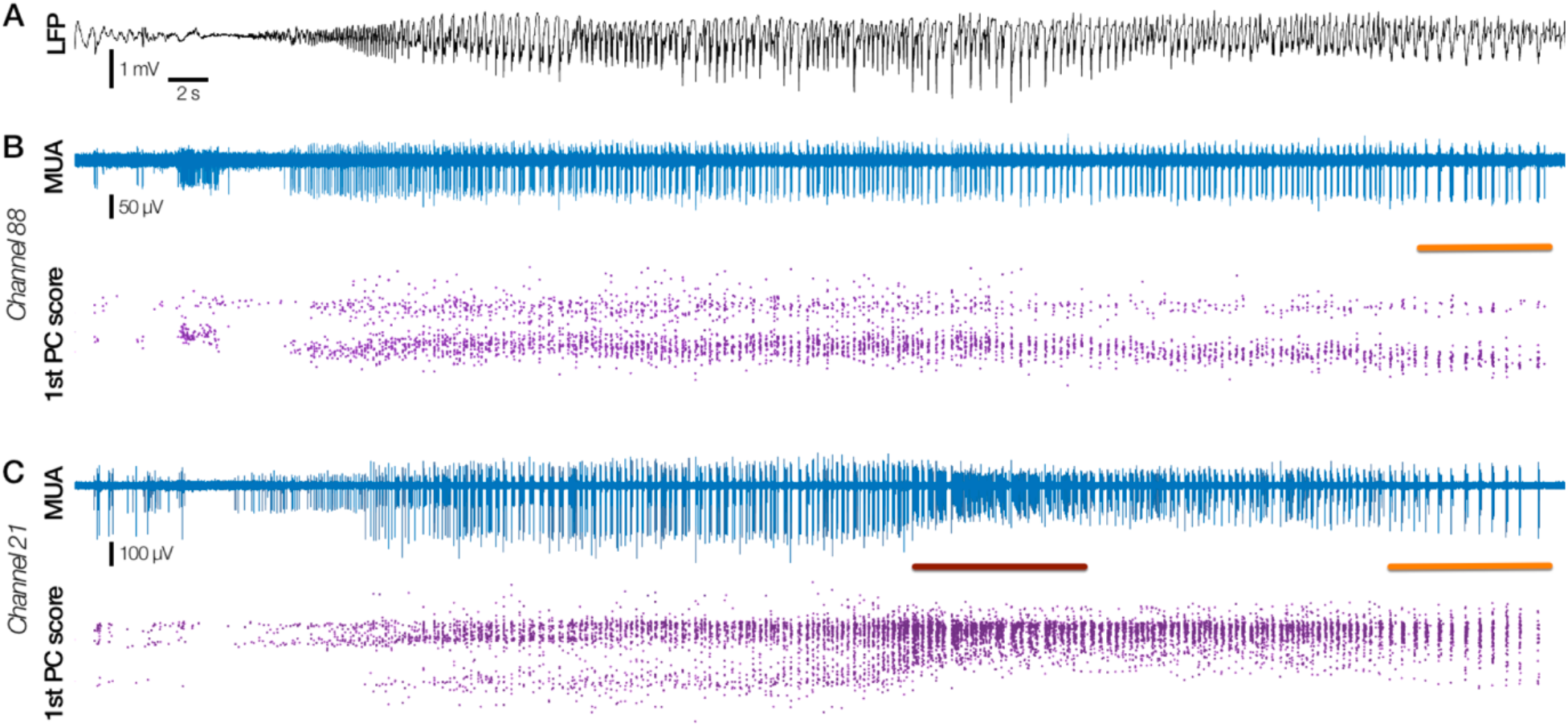
Simultaneous recruitment and penumbral recording. Two types of activity pattern recorded simultaneously in a patient with the Utah array at the edge of the clinically defined seizure spread (Patient 6). **A.** LFP from the closest macro-electrode to the UMA, with time-locked MUA (blue) and first principal component score (purple) through time from channels 88 and 21 in **B** & **C** respectively. Note the stability of waveform and principal component score throughout the seizure in channel 88, with no evidence of tonic firing, while there is a large spike shape change at the same time in channel 21, at the moment of tonic firing (maroon bar, C). Paired orange bars in B and C denote burst firing at the end of the seizure in both locations. These dual activity types both occurred immediately next to the LFP in A, and thus these patterns cannot be differentiated at the macro LFP level.

**Table 3.** Single unit data from Utah array population analyses. Total number of units found in each patient and seizure for population analyses in Utah array cases via the convex hull template-matching method, along with the number of units showing significant increases in firing rate, spike full-width-at-half-maximum (FWHM), and decreases in amplitude during the seizure.

| Patient | Seizure # | Recruited? | Units from convex hull [n] | Units showing FR increase in ictal wavefront [n (%)] | Units showing significant FWHM increase [n (%)] | Units showing significant amplitude decrease [n (%)] |
| --- | --- | --- | --- | --- | --- | --- |
| 1 | 1 | X | 52 | N/A | 2 (4%) | 3 (6%) |
| 2 | 1 | X | 55 | N/A | 9 (16%) | 8 (15%) |
|  | 2 | X | 49 | N/A | 3 (6%) | 2 (4%) |
| 3 | 1 | ✓ | 80 | 77 (96%) | 63 (79%) | 20 (25%) |
|  | 2 | ✓ | 80 | 70 (88%) | 68 (85%) | 38 (48%) |
|  | 3 | ✓ | 79 | 67 (85%) | 66 (84%) | 40 (51%) |
| 4 | 1 | ✓ | 101 | 77 (76%) | 93 (92%) | 45 (45%) |
| 5 | 1 | ✓ | 117 | 82 (70%) | 114 (97%) | 103 (88%) |
|  | 2 | ✓ | 104 | 73 (70%) | 85 (82%) | 63 (61%) |
| 6 | 1 | X | 221 | N/A | 105 (48%) | 86 (39%) |

Similarly, units in recruited cortex showed a significant decrease in spike amplitude during the seizure (Fig. 6C; pre-ictal vs. ictal mean ± SD: 48.82 ± 30.91 µV vs. 34.96 ± 19.54 µV), while penumbral recordings maintained their peri-ictal amplitude (Fig. 6D; pre-ictal vs. ictal mean ± SD: 46.69 ± 16.59 µV vs. 45.08 ± 15.05 µV in fully penumbral cases; 47.00 ± 21.18 µV vs 45.57 ± 20.93 µV in the semi-recruited UMA). The amplitude reduction in recruited tissue was significantly greater than in penumbral/edge case recordings (p < 0.001; one-tailed Mann-Whitney U test), with recruited recordings showing significant (*p* < 0.05) decreases in amplitude during the seizure in 49.3% of units versus 6.4% of units in the penumbra and 38.9% in semi-recruited tissue (Holm-Bonferroni corrected Mann-Whitney U test).

### Spike shape changes in deep structures

To assess the spatiotemporal relationship between waveform alterations and seizure recruitment within patients, beyond the capabilities of the 4 mm^2^ Utah array, we analysed BF array recordings with the equivalent template matching, blind to the clinically-defined seizure onset zone and areas of propagation. In BF array recordings, 120 of 132 units were successfully tracked using these methods. Thirty of 120 single units (25.0%) showed increases beyond a cut- off significance level of *p* < 0.05 (Holm-Bonferroni corrected Mann-Whitney U test) in FWHM (17 seizures from 13 patients), and 30 units (25.0%; 16 seizures from 11 patients) showed reduction in spike amplitude below the same significance cut-off (*p* < 0.05; Table 2). In 9 seizures from 6 patients, single units were simultaneously present on multiple separate BF arrays (on different bundles of microwires, as opposed to different microwires within a single BF), of which 7 seizures (6 patients) showed waveform alterations in at least one unit. Of these, 2 seizures (2 patients) showed significant waveform alterations in dual locations (Patients 14 and 15; Fig. 8; Table 2), while 5 seizures (4 patients) showed both activity types simultaneously (Patients 11, 12, 16 and 21; Video 1).

**Table 2.** Demographics and results for patients implanted with Behnke-Fried arrays. Individual patient demographics for stereo-EEG/Behnke-Fried array cases, including the number of isolated single units found for each seizure in each patient, implant locations that showed significant waveshape alterations, the delay until waveforms surpassed > 1 SD beyond the preictal mean, and whether clinical observations matched these findings. FTBTC: focal to bilateral tonic-clonic; FA: focal aware; FIA: focal with impaired awareness; HC: hippocampus; MC: mid-cingulate; “Spread”: matches clinical observations of seizure spread; ✔+: true positive match for clinical observations; ✔–: true negative match for clinical observations; ✗+: false positive match for clinical observations.

| Patient | Demographics | Implant locations with units | Seizure type | Seizure onset | Seizures analyzed | # units | Microwires with waveform alterations | Time of recruitment (s) | Clinical correlation |
| --- | --- | --- | --- | --- | --- | --- | --- | --- | --- |
| 7 | 29 F | Mesial temporal (Left hippocampus) | FTBTC | Left lateral frontal | 2 | [2, 2] | Left hippocampus | 16.3 | Spread |
| 8 | 29 F | Mesial temporal (Left hippocampus) | FA | Left mesial temporal | 2 | [2, 1] | Left hippocampal head | 16.1 | ✓+ |
| 9 | 55 F | Mesial temporal (Right hippocampus) | FIA | Right lateral temporal | 1 | 9 | — | — | ✓— |
| 10 | 25 M | Mesial temporal (Left hippocampal body) | FIA | Right mesial temporal and right parietal (dual) | 1 | 1 | Left hippocampal body | 42.7 | Spread |
| 11 | 19 M | Frontal (Right anterior cingulate) | FIA | Right lateral temporal | 1 | 3 | Right supplementary motor area | 9.1 | Spread |
| 12 | 20 F | Frontal (Left anterior & mid-cingulate) | FA | Left mesial temporal | 1 | 15 | Left mid-cingulate | 19.5 | Spread |
| 13 | 21 F | Mesial temporal (Left hippocampus) | FA | Left anterior temporal pole | 1 | 1 | — | — | ✓— |
| 14 | 40 M | Mesial temporal/frontal (Left hippocampus & anterior cingulate) | FIA | Left lateral frontal | 2 | [9, 10] | Left hippocampus; Left mid-cingulate | 39.3 (HC); 29.0 (MC) | Spread |
| 15 | 25 F | Mesial temporal (Right hippocampus body & head) | FIA | Right lateral temporal spreading to mesial temporal | 1 | 11 | Right hippocampal body; Right hippocampal head | 9.1 (HCB); 8.0 (HCH) | ✓+ |
| 16 | 35 F | Mesial temporal (Bilateral hippocampus) | FIA | Left mesial temporal | 1 | 5 | Left hippocampal head | 8.0 | ✓+ |
| 17 | 40 M | Mesial temporal (Left hippocampus) | FIA | Left mesial parietal | 1 | 2 | — | — | ✓— |
| 18 | 39 M | Frontal (Right mesial cingulate) | FIA; FTBTC | Right mesial temporal and left orbitofrontal (2 types) | 2 | [8, 4] | — | — | ✓— |
| 19 | 30 M | Frontal (Right mesial cingulate) | FA; FIA | Left mesial temporal and left insula | 2 | [1, 1] | — | — | ✓— |
| 20 | 23 F | Mesial temporal (Right hippocampus) | FIA | Right posterior temporal | 1 | 3 | — | — | ✓— |
| 21 | 35 M | Mesial temporal (Bilateral hippocampus) | FA | Right insula and somatosensory cortex | 3 | [7, 8, 7] | Right hippocampal head | Sub-threshold | Spread /X+ |
| 22 | 44 F | Frontal (Right anterior cingulate) | FTBTC | Left cingulate | 1 | 1 | Right anterior/mid-cingulate | 15.2 | Spread |
| 23 | 51 F | Frontal (Left anterior cingulate) | FIA | Left mesial temporal | 1 | 4 | Left anterior cingulate | 12.6 | Spread |
| 24 | 20 M | Frontal (Left anterior cingulate) | FIA | Left posterior temporal | 1 | 1 | — | — | ✓— |
| 25 | 30 F | Frontal (Right anterior cingulate) | FIA | Left mesial temporal | 1 | 1 | — | — | ✓— |
| 26 | 20 M | Mesial temporal (Right hippocampus) | FIA | Right mesial temporal | 2 | [1, 2] | Right hippocampal body | 10.4 | ✓+ |
| 27 | 32 M | Mesial temporal (Bilateral hippocampus) | FIA | Bilateral mesial temporal | 3 | [4, 3, 3] | Hippocampal body | 9.8 | ✓+ |

We found significant waveform alterations in 13 patients. Five of these had a clinically defined SOZ in the mesial temporal lobe, and simultaneously recorded single units in the ipsilateral hippocampus demonstrated significant waveform alterations (Patients 8, 15, 16, 26 & 27; Table 2). In 7 further patients, significant waveform alterations were found in tissue consistent with putative seizure spread due to proximity to the SOZ, or due to seizure generalization. In one case (Patient 10) we found waveform alterations in the contralateral hippocampal body, consistent with propagation of the seizure through the hippocampal commissural fibers. Conversely, in the 8 patients showing no significant waveform alterations, the clinically defined SOZ was anatomically distant in all cases (Patients 9, 13, 17–20, 24 & 25; Table 2).

To assess whether the classification of ictal recruitment via waveform alterations was consistent with the clinically defined SOZ and regions of spread, the time from earliest ictal activity to a consistent (≥ 1 s duration) increase in FWHM ≥ 1 SD above the mean preictal level for each unit was calculated. FWHM was used independently of amplitude for these tests to control for any potential fluctuations in amplitude introduced by the reference electrode during seizures. Recordings determined to be in the SOZ showed a mean (± SD) delay of 10.23 ± 3.03 s (*n* = 6 seizures from 5 patients), while those deemed to be in regions of spread showed a mean (± SD) delay of 22.96 ± 12.59 s (*n* = 8 seizures, 8 patients; *p* < 0.05, Mann-Whitney U test). In one case, single unit waveforms remained stable throughout a focal to bilateral tonic-clonic seizure (Patient 18; Table 2), further countering the possibility of transient movement artefact introducing instability in the waveforms.

In one instance of a patient recorded with BF arrays in the hippocampal head and body, a peculiarly discrete unit cluster due to a very large amplitude spike (mean = 354 µV; background noise level = 25 µV) enabled us to follow its action potential through the ictal transition without the need for template matching via convex hulls, despite marked changes to spike shape and increases in other unit activity (Fig. 8; Video 2; Schevon *et al*., 2019). In this example, the seizure initiated very close to, but not at the electrode site. The ictal wavefront arrived at the electrode approximately 8 s after seizure initiation (Patient 15; Table 2). The action potential amplitude was stable during both the pre-ictal period and the moments after seizure initiation, but reduced sharply upon the abrupt increase in firing rate indicating ictal recruitment (Fig. 8A & B, magenta dashed line; preictal vs. ictal mean ± SD: 354.2 µV ± 45.7 µV vs. 265.7 µV ± 33.4 µV; *p* < 0.001, Mann-Whitney U test). The inter-spike interval to spike amplitude relationship was similar to animal models of recruitment (c.f. Supplementary Fig. 4 in Merricks *et al*., 2015; 0 Mg^2+^ *in vitro* cell-attached mouse slice model), and echoed the template matched results in the Utah array (note the similarity between Fig. 4B and Fig. 8D).

**Fig. 8.**
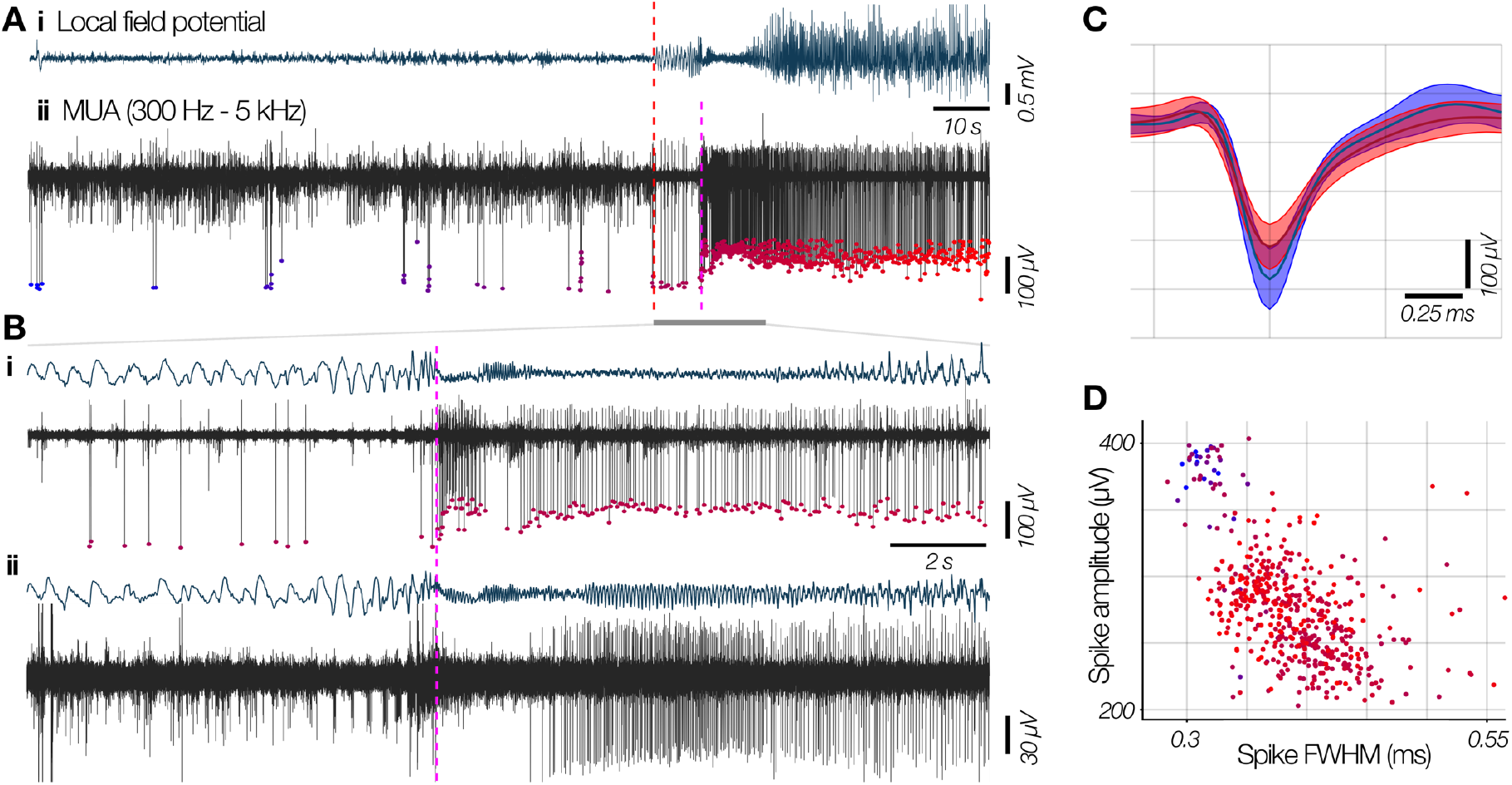
Ictal recruitment in the mesial temporal lobe recorded with BF electrodes. **A. (i)** Broadband LFP from the closest macro contact to the microwires in a BF electrode in the mesial temporal lobe during a spontaneous seizure, and **(ii)** the bandpass filtered MUA from one of the microwires in the temporal pole. “Global” seizure onset is denoted by the red dashed line, with subsequent local recruitment to the seizure at the pink dashed line. **B.** Magnification of the region denoted by the grey bar in A, showing seizure onset through passage of the ictal wavefront in LFP and MUA (colours maintained) in the same microwire **(i)**, and a microwire from a nearby separate BF in the hippocampal body **(ii)**. Note the pre-recruitment stability in the spike amplitude, which is immediately reduced upon ictal invasion, and the simultaneous quiescence in the hippocampal body, followed by a similar, time-delayed amplitude change after recruitment. **C.** Mean ± SD of waveforms prior to ictal invasion (blue) and after recruitment during the seizure (red), showing reduction in amplitude and increase in FWHM in A(ii). **D.** Spike amplitude versus spike FWHM for the defined unit, with color maintained from A, transitioning from blue to red through seizure invasion. Note the bimodal clusters that correspond to pre- and post-recruitment, and the similarity to Fig. 4B.

Notably, a separate single unit recorded at an adjacent BF array showed spike shape changes which developed 1 second later, consistent with seizure propagation (Fig. 8B ii; Table 2). The distinct time course of these two units recorded in the hippocampal head and body show that the alterations are unit-specific and not caused by local changes as a result of the seizure or by movement artefacts (the seizure had a relatively calm semiology, without stressing the recording device).

Simultaneously, action potential FWHM was stable prior to ictal recruitment, increasing significantly after the passage of the ictal wavefront (preictal vs. ictal mean ± SD: 0.33 ms ± 0.04 ms vs. 0.40 ms ± 0.05 ms; *p* < 0.001, Mann-Whitney U test). The time course of this transition followed the same progression as the spike amplitude (Fig. 8D), as can be seen in the pre-recruitment versus post-recruitment mean (± SD) waveforms (Fig. 8C, blue and red respectively).

### Template matching accuracy

Spike matches from the convex hull method were found to be significantly more likely to arise from their assigned peri-ictal units than by chance, as calculated by comparing each matched waveform’s similarity to its peri-ictal unit’s distribution of voltages through time (Fig. 3; “Spike metrics” in Methods). The “null” distribution for expected match probabilities by chance was calculated through comparing each waveform’s similarity to the peri-ictal voltage-time distributions of all other units. Comparing intra- to inter-unit similarities in this way found a significantly higher similarity between template matched units and their presumed peri-ictal unit than to the “null” distribution of matches to other units (*p* < 0.001; Mann-Whitney U test; Fig. 3B).

To confirm that these findings were not a result of the template matching method introducing an unknown variable that affects these measurements, results from the original spike sorted data and those originating from the convex hull method on the pre-ictal period were compared, finding little difference between the traditional cluster cutting results and the convex hull matched results (Fig. 6 A-D i, inset cumulative histograms).

### Neuronal firing rates through the ictal transition

In animal models, it is possible to isolate single units experimentally, by visually guiding electrodes directly onto cells (Trevelyan *et al*., 2006). This obviously is not possible in human recordings, and consequently, there is a dearth of evidence about the firing patterns of human neurons through seizures. Instead, previous studies have identified the ictal wavefront in terms of multi-unit activity (Schevon *et al*., 2012; Smith *et al*., 2016). We therefore analysed the firing rates of template matched unit populations throughout ictal activity and related these to the ictal wavefront (UMA recordings showing tonic to clonic firing; Patients 3–5; see Methods). Single unit firing rate increased during ictal recruitment in all seizures with 446 (79.5%) of 561 units showing greater than 3 SD increase in firing rate, and only 1 unit in the entire population showing a greater than 3 SD decrease in firing rate (range of single units with > 3 SD increase per seizure: 70% – 96%; see Table 3). An example seizure demonstrating these trends is shown in Fig. 9.

## Discussion

The analyses presented here explored the impact of spike shape alterations in spontaneously occurring seizures, in humans, how these alterations relate to the underlying ictal territories, and whether the ictal wavefront – the source of seizure propagation – involves local neuronal firing or is dominated by subthreshold or synaptic activity. We have previously shown that traditional spike sorting methods fail upon ictal invasion of the recording site (Merricks *et al*., 2015), however, in these recordings it was not possible to differentiate between loss of spike sorting ability due to intrinsic waveform alterations, or due to hypersynchronous activity (Trevelyan *et al*., 2006, 2007; Schevon *et al*., 2012; Weiss *et al*., 2013). Here, we sought to overcome this limitation through novel methods to track units through the ictal transition, retaining the identities of putative individual neurons. We hypothesized that the temporary loss of clusters was due in part to transient alterations to spike shapes, as opposed solely to obfuscation of stable spike shapes by the sudden increase in activity in the MUA, and that units at brain sites not demonstrating evidence of ictal recruitment via tonic firing in the MUA would remain stable.

We applied this method in two types of microelectrode recordings: UMA and BF arrays, representing neocortical and deep structure, particularly hippocampal, seizure foci. Although there must be some expected loss of detection sensitivity due to interference from highly synchronous firing along with the reduction of amplitude of some units below the noise threshold, the method proved remarkably effective, enabling us to define firing rate metrics for the majority of units throughout the seizure, highlighting a lack of neuronal quiescence during seizures.

In both types of recording, template matched units during the ictal period displayed two types of activity: deformation of waveshapes across the population, or largely stable waveforms. The UMA afforded the ability to detect local ictal recruitment through characteristic MUA firing, and these types of activity corresponded to recruitment and penumbral recordings respectively (Fig. 6). Waveform alterations recovered after seizure termination and showed a stereotyped response across seizures, highlighting that a single neuron’s wave shape change, in response to the synaptic barrage of upstream ictal activity, is maintained across multiple seizures hours apart. Note that the template matching method would equally favor alterations to waveform in any dimension, and so if these changes were purely a result of the methodology, we would reasonably expect the template to capture spikes with larger amplitude and decreased FWHM at an equal rate. In such a case, we would anticipate a broadening of the distribution of these features during the ictal period. Instead, we see a clear shift in the distributions to the right and left in the FWHM and amplitude respectively, and found no evidence of increased spike amplitude during seizures, arguing for a consistent physiological cause (Fig. 6A&C).

Detection of the ictal wavefront was not possible in BF arrays, due to a combination of lower neuronal density in mesial structures relative to layer 4/5 neocortex (Pakkenberg & Gundersen, 1997; Keller *et al*., 2018), and a reduced “listening sphere” relative to UMAs due to higher impedance (BF: 50-500 kΩ; UMA: 80-150 kΩ; Tóth *et al*., 2016). Nonetheless, the same waveform features were detected, and their presence correlated well with the clinical assessment of SOZ and seizure spread (Table 2). Furthermore, BF arrays allowed for sampling of multiple sites in a given patient, and the timing of waveform alterations correlated well with clinical observations, with seizure spread locations showing delayed waveform changes compared to recordings from SOZ regions (22.96 vs. 10.23 s respectively). The longest delay until recruitment occurred after spread to the contralateral hippocampus (Patient 10; 42.7 s). The only case with discordance between single unit and clinical data showed a statistically significant increase in FWHM during the seizure, but at no point did the mean FWHM surpass the threshold of 1 SD above the preictal mean (Patient 21; Table 2). In this instance, the clinical SOZ was in the right insula and somatosensory cortex, with waveform alterations in the right hippocampus. While recruitment of the hippocampus is plausible, this may represent a false positive as a result of the temporally coarse statistical test.

The UMA and BF array recordings together provide evidence supporting the dual-territory model of seizures: an ictal core with waveform changes, coexisting with a penumbral territory with stable spike shapes. Combined, these indicate that the definitions of ictal recruitment and penumbra are maintained at the level of single neurons, and waveshape change can be considered a defining feature of recruitment to the seizure. In a subset of recordings, stable action potentials were found simultaneously with waveform changes both in separate BF arrays (5 seizures) and on the same UMA (1 seizure; Patient 6), with clinical correlation matching these observations in all cases. In the UMA, clinical observations were consistent with location of the UMA at the outer boundary of seizure invasion, and thus we posit this is a simultaneous recording of both recruited and penumbral cortex (Fig. 7).

Furthermore, in 2 seizures, recruitment was found at multiple BF arrays, consistent with clinically defined SOZ and regions of spread, along with relative delays in keeping with anatomical distance (Fig. 8; Patients 14 & 15; mid-cingulate to hippocampus in 10.3 s, hippocampal head to body in 1.1 s respectively). Note also, that in Patient 6 there is an increase in waveforms of brief duration during the ictal period (data available online; population data shown in Fig. 6B). Given the UMA’s proximity to the oncoming ictal wavefront, this increase may be explained by an increase in firing of fast-spiking interneurons, which have been shown to exhibit action potentials of shorter duration (McCormick & Feeser, 1990; Csicsvari *et al*., 1999; Peyrache *et al*., 2012) and would corroborate the penumbral feedforward inhibition model (Trevelyan *et al*., 2007; Cammarota *et al*., 2013; Parrish *et al*., 2019).

More specifically, the waveform changes found in recruited tissue are in keeping with those observed in animal models, and are indicative of the shortening and broadening of action potentials associated with PDS (Traub & Wong, 1982). This was especially evident in a unique BF array recording wherein a unit was able to be tracked without need for extra methods due to its amplitude being 14 times that of the background noise (Patient 15; Fig. 8), with feature alterations strikingly reminiscent of the UMA population data (c.f. Figs. 4B & 8D), with timing in keeping with recruitment, during tonic firing (c.f. Figs. 7 & 8).

Even so, the extent to which these waveforms alter is likely underrepresented in the population data, due to changing detection sensitivity from interference between synchronous spikes or the reduction of amplitude of some spikes below the noise threshold. The template-matching method was designed to minimize false negatives in order to capture as much single unit activity during seizures as possible, but it is likely that spikes undergo large enough changes to be lost outside the convex hull, or below threshold. Indeed, even physiological bursting has been shown to result in substantial alterations to extracellularly recorded action potentials (Harris *et al*., 2000; Henze *et al*., 2000). As such, these results are necessarily a snapshot of the total activity of any individual neuron during the seizure, and yet still show significant changes to waveform.

Finally, our data also demonstrate that the passage of the ictal wavefront is marked by tonic, local neuronal firing (Fig. 9), as opposed to the wavefront being a signature of increased K^+^ concentration or similar subthreshold phenomena. Studies of single unit activity recorded in humans have rarely reported such a finding. Although placement of the electrodes likely plays a large role, our findings in this paper suggest that extreme, rapid waveform alterations may obscure the presence of the wavefront when standard spike sorting methods are used. Note that these firing rate statistics are probabilistic, having been scaled by the probability each spike was a match to its preictal unit (see Methods), and as such are conservative and thus the increase cannot be attributed purely to increases in background spikes being matched by the convex hull method.

**Fig. 9.**
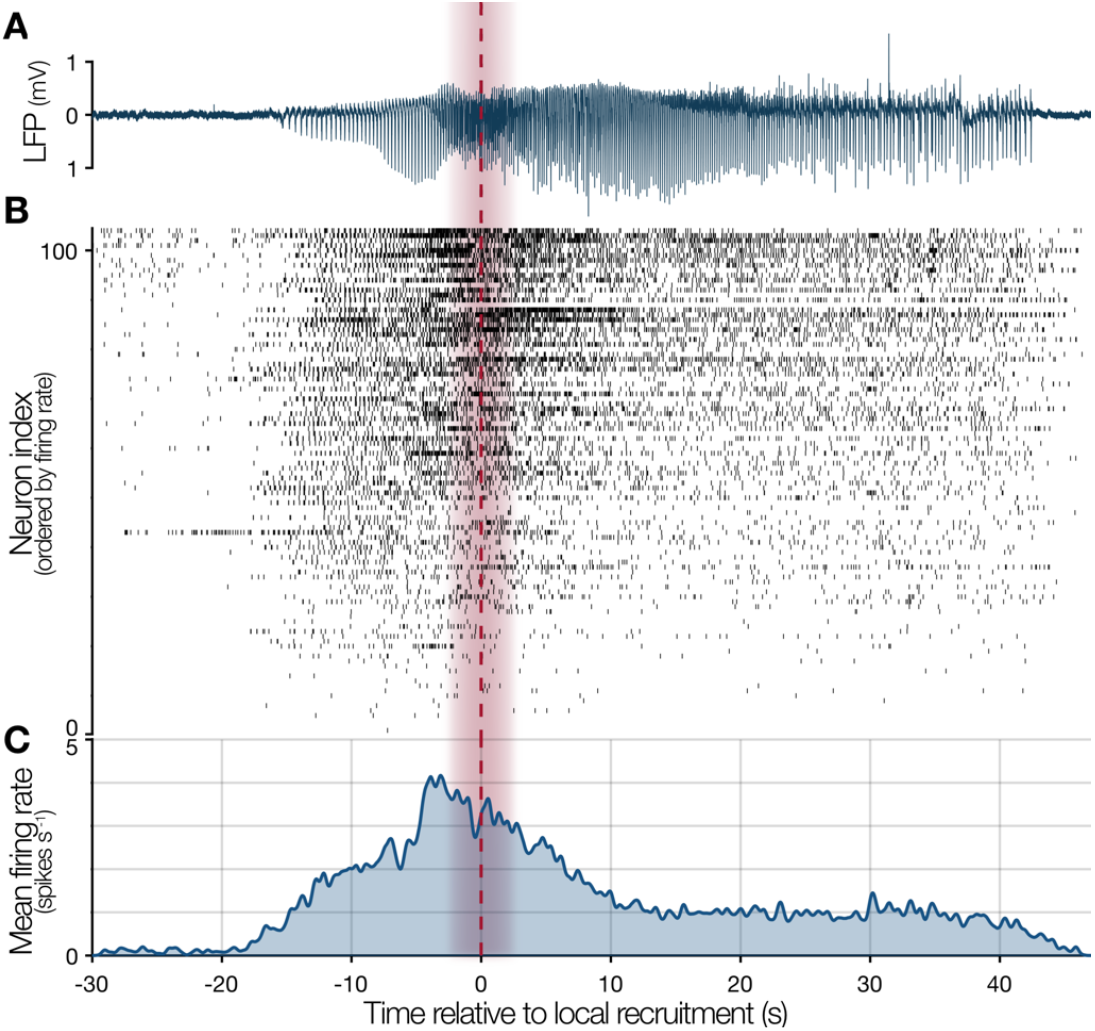
Neuronal firing through ictal recruitment in an example focal seizure. **A.** Local field potential (LFP) from an example channel during seizure 1 in Patient 3, with the calculated passage of ictal wavefront marked by red dashed line, with ± 2 seconds shaded red. **B.** Raster plot of all units found using convex hull template matching in this seizure, ordered by firing rate, and plotted relative to the moment of ictal recruitment at that channel. **C.** Probabilistic instantaneous firing rate of the population of single units as estimated by convolving a Gaussian kernel (200 ms SD) over the spike times. The firing rate shown is probabilistic by scaling the Gaussian kernel’s amplitude by the likelihood of each individual spike originating from its assigned preictal unit, as calculated by its voltage-time probabilities (see Methods & Fig. 3). As such, the firing rate has not been biased by excessive matching of dissimilar waveforms during the ictal activity. Note the intense, tonic firing during the seizure invasion, and sustained, above baseline firing until seizure termination.

This method of tracking single units across the ictal transition will enable the use of both human and animal recordings to address open questions regarding the mechanism of seizure spread. An immediate application, given the neuronal activity presented here, is to study how cell types relate to the propagation of pathological activity, with considerable debate having focused recently on the role of interneurons in seizure spread (Grasse *et al*., 2013; Elahian *et al*., 2018; Magloire *et al*., 2018; Miri *et al*., 2018; Weiss *et al*., 2018). All studies to date, to our knowledge, have assessed cell-type specific activity at seizure onset in regions without waveform alterations, and thus we suggest are recordings from penumbral territories. As such, we anticipate elucidation of these mechanisms will come from population data confirmed to be in recruited tissue. These methods lay important groundwork for analyses into how ictal propagation relates to the underlying firing of local inhibitory and excitatory cells.

## Video Legends

**Video 1. Location specific waveform changes during a spontaneous human seizure**

Local field potential (top) and spike amplitudes for three units in three different locations (middle; red, blue and purple), with associated spike waveforms shown to the right. Shading shows the mean ± 2 SD for these units’ preictal spike amplitudes. The locations of the BF microwires that recorded these units are shown below, with colors maintained. Note the three activity patterns: cessation of firing at seizure onset in the anterior temporal lobe (blue), stability followed by loss of spike amplitude in the anterior cingulate (red), and stability throughout in the mid-cingulate (purple).

**Video 2. Single unit waveform alterations in an ictal Behnke-Fried recording**

Single units undergo waveshape changes upon recruitment to the seizure, shown in real-time. Upper trace: MUA bandpassed signal (300 Hz to 5 kHz; white), with current time shown in red, earliest electrographic ictal activity in the patient occurs at dashed magenta line, with local recruitment occurring at blue dashed line. Lower panels: shaded regions show mean ± 2 SD for preictal single units in red and blue, and lower amplitude multiunit activity in yellow (left). Waveforms are displayed in real-time, with colors matching their assigned unit, and color saturation showing the probability of a true match to that unit. The first two principal component scores for these waveforms are shown on the right, with colors maintained. Note the stability of waveforms prior to local recruitment, including after seizure onset, followed by marked loss of amplitude at the moment of recruitment with associated tonic firing.

