## Supplementary figures and images for "Neuronal firing and waveform alterations through ictal recruitment in humans"

### Video 1

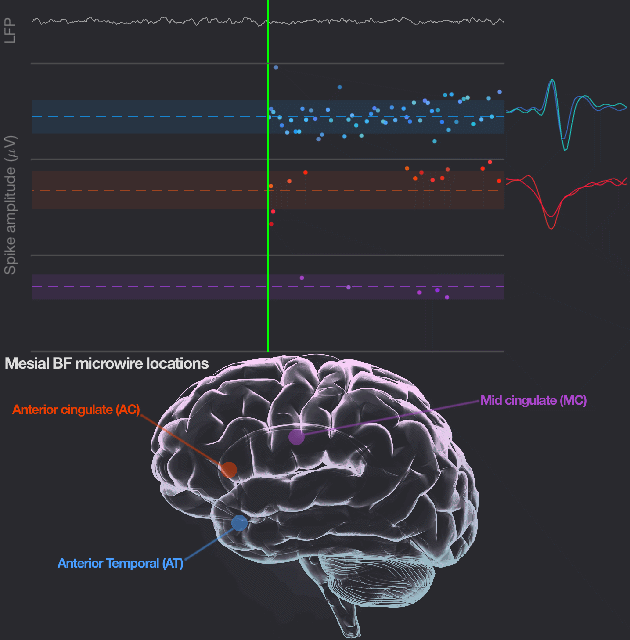
